# Computational modeling of necrosis in neural organoids

**DOI:** 10.1101/2025.04.30.651571

**Authors:** Aishwarya Pantula, Bo Zhou, Itzy E. Morales Pantoja, Alisa Fedotova, Derosh George, Dowlette-Mary Alam El Din, Alexandra Lysinger, Lena Smirnova, David H. Gracias

**Affiliations:** Department of Chemical and Biomolecular Engineering, Johns Hopkins University, Baltimore, MD 21218, USA; Department of Biomedical Engineering, Johns Hopkins University, Baltimore, MD 21218, USA; Center for Alternatives to Animal Testing, Department of Environmental Health and Engineering, Bloomberg School of Public Health and Whiting School of Engineering, Johns Hopkins University, Baltimore, MD 21205, USA; Department of Materials Science and Engineering, Johns Hopkins University, Baltimore, MD 21218, USA; Department of Chemistry, Johns Hopkins University, Baltimore, MD 21218, USA; Sidney Kimmel Comprehensive Cancer Center, Johns Hopkins School of Medicine, Baltimore, MD 21205, USA; Laboratory for Computational Sensing and Robotics (LCSR), Johns Hopkins University, Baltimore, MD 21218, USA; Department of Oncology, Johns Hopkins University School of Medicine, Baltimore, MD 21205, USA; Institute for NanoBioTechnology, Johns Hopkins University, Baltimore, MD 21218, USA; Center for MicroPhysiological Systems (MPS), Johns Hopkins School of Medicine, Baltimore, MD 21205, USA

## Abstract

Neural organoids (NOs), also known as brain organoids, are derived from human-induced pluripotent stem cells and are Microphysiological Systems (MPS) of the brain that can recapitulate key aspects of neurodevelopment. They enable *in vitro* studies of brain development and disease mechanisms, providing disease models for various neurodegenerative or neurodevelopmental/degenerative disorders like Alzheimer’s disease, microcephaly, and autism. There are many protocols to generate NOs with different complexities and sizes, varying from 400 μm to several mm in diameter, with a starvation-induced necrotic core eventually forming depending on the diameter and culture conditions. Thus, they can benefit from vascularization and more optimal culture conditions. There have been several attempts to decrease necrosis while growing larger NOs, such as by using orbital shaking or 2D/3D microfluidic chips, but only with limited success. In this study, we describe a 3D finite element model to simulate O_2_ starvation-induced necrosis in NOs using the Damköhler Number (*Da*) and the Michaelis-Menten kinetics. We measured the necrotic areas in NOs using fluorescent imaging and used them to calibrate the model with a specific *Da*. Using these calibrated values, we systematically compared simulations of different NO culture methods—static, orbital shaking, and microfluidic flow around organoids—highlighting their relative impacts on nutrient diffusion and necrosis. We observed that these culture strategies cannot prevent necrosis beyond a diameter of ∼800 μm. Based on these findings, we propose that 3D spatial perfusion, achieved through uniformly distributed fluidic capillaries within the NO, could significantly reduce necrosis. We conducted parametric studies on capillary spacing, density, and layout. Our calibrated model offers insights for designing next-generation microfabricated bioreactors and culture devices, not just for NOs but also for all 3D tissue engineering and organoid research.

## 1. Introduction

Cell culture has progressed from planar to three dimensional (3D) cell culture models. Concurrent advances in stem-cell biology have led to organoid models— micron-sized organ-specific 3D tissues grown from pluripotent stem cells (induced and embryonic) *in vitro*. These constructs mimic organotypic structure and function, and are used to build Microphysiological Systems (MPS) used to model various tissues (e.g., brain, intestine, liver, pancreas). MPS models are essential in biomedical research for studying organ development, disease, and drug testing.^1–3^

Neural organoids (NOs) or brain organoids, derived from human pluripotent stem cells, represent a significant breakthrough in modeling the human brain.^4–8^ These self-organizing 3D structures replicate key cellular compositions and architecture, forming distinct brain-like regions with neurons, astrocytes, and oligodendrocytes. NOs are invaluable tools for understanding early brain development, studying neurodevelopmental and neurological disorders (e.g., microcephaly, Autism Spectrum Disorder (ASD), Alzheimer’s disease (AD)), and advancing drug discovery. NOs offer more physiologically relevant and ethical alternatives to animal models.^4–8^ However, NOs and organoids in general are still simplified models, which currently lack the functional and spatiotemporal organization observed in real organs. Rather, they mimic a functional unit of an organ. ^9^ While it is clear that increasing organoid size can enhance functionality, significant biological and technical challenges limit this approach. In particular, nutrient and waste diffusion limit the growth of organoids in culture, especially beyond diffusion scales. ^10–13,14^ In particular, oxygen (O_2_) transport can be used to capture the necrotic effects. O_2_ is the critical determinant of necrosis over glucose because of its lower concentration in organoid media and higher consumption rate (1–2 orders of magnitude less), despite it having a diffusivity *∼*10 times higher.^15^ The thickness of cultures and even tissue slices is less than the diffusion limit of O_2_ in them (below 200-300 μm). (**Figure 1A**). In contrast, in 3D cultures like NOs, the lack of vascularization causes O_2_ deprivation in the densely packed core, resulting in a dead core or necrosis.^16–18^ (**Figure 1B**).

**Figure 1.**
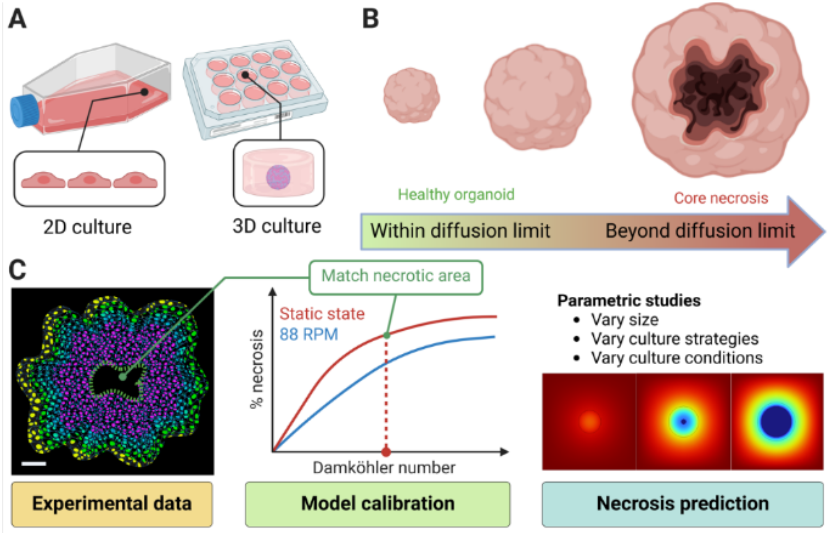
Modeling necrosis in 3D cultures: **(A)** Schematic illustrating 2D vs 3D tissue cultures. **(B)** Schematic diagram showing that while necrosis is not a concern for smaller organoids, it occurs in the dense core regions of larger organoids. **(C)** Workflow of our overall strategy for computational modeling of necrosis within 3D culture using NOs. We identify necrosis in stained slices of NOs (*left*) and fit the percentage (%) of necrotic area to get the respective Damköhler number (*Da*) (*middle*) and proceed with parametric studies for the NO in our 3D CAD models (*right*).

Indeed, researchers have been exploring various approaches to minimizing necrosis over a traditional static culture. For example, NOs have been co-cultured with endothelial cells or xenoimplanted in a mouse with limited permeation of the vasculature.^19–21^ To enhance media perfusion within NOs, there have been attempts to enhance transport using forced convection, for example, by shaking on an orbital platform.^13,22– 24,20,25,28,29^ Several studies have investigated the flow of media around NOs using microfluidic chips or micropillars.^20,25–27^ However, most of these dynamic culture approaches have been shown to work for NOs with diameters around 600 µm. Previous studies have utilized finite element modeling (FEM) to explore O_2_ consumption and hypoxia-induced necrosis in a variety of cell cultures, including NOs, examining the effects of culture strategies like shaking and microfluidic systems on O_2_ transport and viability^14-16,38,39^. These efforts, often validated with experimental data, have applied convection-diffusion equations coupled with Michaelis-Menten kinetics to model O_2_ diffusion and consumption. Due to the computational cost of the complex geometries and meshing, most of these models are 2D, often focused on one culture strategy with limited modeling of dynamic culture systems like shaking, spinning, or agitation. ^14-16^ These studies typically examine smaller organoids (600–800 μm) and do not extend to larger constructs exceeding 1–2 mm in diameter, where diffusion limitations become more critical. Consequently, there is a lack of systematic, comparative modeling to evaluate how necrosis progresses across different culture strategies, especially for increasing organoid size (beyond 1000 μm) and metabolic demand. Hence, addressing this gap requires a unified modeling framework capable of predicting and comparing necrotic outcomes across commonly used methods, ultimately guiding the design of more effective, scalable organoid culture systems.

We developed a 3D computational model (COMSOL Multiphysics) to investigate the progression of necrosis in NO culture. Our model uses 3D CAD geometries built in Solidworks (Dassault Systèmes) and use a convection-diffusion-reaction Multiphysics interface. Since organoids may differ in cellular composition from culture to culture, the metabolic consumption rates in the NO can vary by two orders of magnitude. ^28,29,32,33^ This variability presents a major challenge for predictive modeling. To address this challenge, we developed a generalized calibration method using the Damköhler number (*Da*). This dimensionless number represents the ratio of reaction to diffusion rate in a system, to capture the competing effects of O_2_ cellular consumption vs. transport in the tissue/culture media.^30,31^

We established calibration curves across a *Da* range of 0.01 to 20 (**Figure 1C**), extending the model beyond our NO system to a wide range of 3D culture platforms for cells, tissues, and other organoids. Furthermore, to account for differences in NO generation protocols,^28-33^, we generated additional calibration curves spanning the same *Da* range across multiple orbital shaking speeds, thereby enhancing the adaptability of our model to diverse dynamic culture conditions (**Figure S8**).To calibrate our simulation model, NOs were cultured from NIBSC8 iPS cells (female line, normal karyotype) for eight weeks under static and orbital shaking (88 RPM) conditions. We then calibrated our FEM model to our NO system by matching up the necrotic area found in both of the conditions to the model using a single *Da*. Using that *Da*, we simulated and compared the necrosis in different modes of NO culture, including static culture, orbital shaking, and flow around NO in a microfluidic chip, with the size of the NO, establishing the size limitation of the efficacy of each culture strategy. Based on these findings, we tested the concept of 3D spatial perfusion within NO (using synthetic capillaries) for optimal O_2_ transport. We investigated the effects of microfluidic flow rate on the necrotic area for designs that flow media around (flow-through microfluidic chip) and within (spatial perfusion) the NO. We demonstrated that by optimizing capillary density, flow systems that enable spatial perfusion within a NO eliminated necrosis even for size scales beyond the diffusion limitation of dynamic cultures. We also demonstrated that by using different capillary layouts, we can enable spatiotemporal patterning alongside spatial perfusion in the culture devices.

## Materials and methods

### iPSC maintenance, NO generation, and cultivation

We differentiated NOs from induced Pluripotent Stem Cell (iPSC) line NIBSC-8 (female origin, UK National Institute for Biological Standards and Control (NIBSC)). The cell line is mycoplasma-free, with normal karyotypes, and was differentiated following our previously published two-step protocol. ^34^ Briefly, we cultured hiPSCs in mTESR-Plus medium (StemCell Technologies at 5% O_2_, 5% CO_2,_ and 37 °C. We differentiated stem cells in a monolayer to neural progenitor cells (NPCs) using a serum-free, chemically defined neural induction medium (Gibco, Thermo Fisher Scientific). We then expanded and assessed the expression of NPC markers, Nestin, Sox2 for quality control. We distributed a single cell suspension of 2 × 10^6^ NPCs per well into uncoated six-well plates to initiate NO differentiation. We kept our cultures at 37 °C, 5% CO_2_, and 20% O_2_ under constant orbital shaking (88 RPM, 19 mm orbit) to form 3D aggregates. After 48 hours, we induced differentiation with serum-free, chemically defined standard differentiation medium: B-27™ Plus kit, 2% Glutamax (Gibco, Thermo Fisher Scientific), 10 ng/mL human recombinant Glial Cell-Derived Neurotrophic Factor (GDNF) (GeminiBio™), 10 ng/mL human recombinant Brain-Derived Neurotrophic Factor (BDNF) (GeminiBio™), 2% Glutamate, and 1% Pen/Strep (Gibco, Thermo Fisher Scientific). We changed approximately 75% of the medium three times a week.

### Culture conditions for NOs

We evaluated two commonly used mechanical conditions to assess their impact on NO culture and quantify necrotic areas. We cultured NOs for eight weeks on an orbital shaker at 88 RPM with a 19 mm orbit and on a static petri-dish. We maintained all cultures at 37 °C, 5% CO2, and normoxic conditions. We considered a sample size of three NOs for model calibration in each culture condition.

### Cryopresectioning

We fixed the NOs in 4% paraformaldehyde for 45 min, followed by incubation in a sucrose gradient (10-30%) and then embedded them in an optimal cutting temperature (OCT) compound. We then sectioned them at 50 µm thickness using a cryostat.

### Immunohistochemistry

Whole organoids or crysectioned slices were permeablilized with 1 mL permeabilization solution (1X PBS, 0.1% Triton X) for 30 min at 4 °C,,blocked with 100% BlockAid™ (Thermo Fisher Scientific) on a shaker for one hour at 4°C. Then, we incubated NOs at 4°C for 48 h with a combination of primary antibodies (**Table S2**:) in 10% BlockAid™, 1% BSA (Bovine Serum Albumin), and 0.1% Triton X in PBS. We washed NOs in PBS three times and incubated with secondary antibodies conjugated with Alexa 488, Alexa 568, or Alexa 647 (1:500, Thermo Fisher Scientific) and Hoechst 33342 trihydrochloride (1:2000, Invitrogen) Organoids/slices were in 10% BlockAid™, 1% BSA, and 0.1% TritonX in PBS for 24 h. We mounted organoids/slices on slides with coverslips and Prolong Gold antifade reagent (Molecular Probes); negative controls were processed omitting the primary antibodies. We took images with a Zeiss UV-LSM 700 confocal microscope. We performed further imaging processing and quantification of the NO area using ImageJ2 (version 2.14.0/1.54f).

### Modeling O_2_ transport and consumption in NO

We developed a 3D computational model for O_2_ transport using the COMSOL Multiphysics 6.1 software (COMSOL AB, Stockholm, Sweden) and the Reaction Engineering add-on module. We simulated four commonly used NO culture strategies. (1) Static state culture, (2) orbital shaking culture, (3) microfluidic flow of media around the NO, and, (4) media flow within NO using 3D spatial perfusion. We modified a general convection-diffusion-reaction model to simulate the O_2_ diffusion and uptake to generate concentration profiles across a cross-section of the NO. We divided our system into discrete components or elements, applied boundary conditions and partial differential equations, and generated solutions for velocity, pressure, and concentration fields over the geometry. **(More details in Section S1)**. Doing this allowed us to represent the solutions of the vector fields (pressure, velocity, or concentration) precisely over more complex geometries over bulk geometry solutions.^35^

### O_2_ diffusion and uptake

We used a diffusion-reaction model using the ‘Transport of dilute species’ module to simulate the static state culture of NOs. We used Fick’s law to describe O_2_ diffusion through media and NOs, while assuming an incompressible flow,

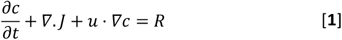

where,

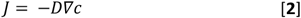

Here, *c* denotes the concentration of the diffusing component (O_2_ in [mol/m^3^], depending on the boundary, *D* is the diffusion coefficient of O_2_ in either media or the NO [mol/m^3^s]. *R*_*i*_ is the rate of O_2_ consumption in NO, *u* is the velocity field [m/s], *∇* the standard vector differential operator (*nabla*) where, 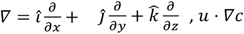 is the advective flux and *∇· J* is the diffusive flux. ^36^

We simulated the O_2_ uptake in the NO using Michaelis-Menten kinetics, coupled with a consumption step down to indicate necrosis once the critical concentration is achieved in the element.

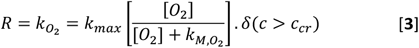

Here, *k*_*max*_ is the maximum O_2_ consumption rate [mol/m^3^s], 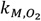 is the Michaelis Menten constant representing the concentration where consumption drops to half its maximum value. *c*_*cr*_ is the critical O_2_ concentration, the O_2_ concentration below which necrosis is assumed to occur due to hypoxia. *δ* is a step-down function that shows the halting of O_2_ consumption due to the cells in the region dying, leading to necrosis.

To create a smooth step down once the critical concentration is reached, we used the Heaviside function with a continuous second derivative (*flc2hs*) function to define *δ·*^37^ We considered estimates of various parameters that were used from previous literature^22,38,39^ and summarized them in **Table 1**.

**Table 1:**
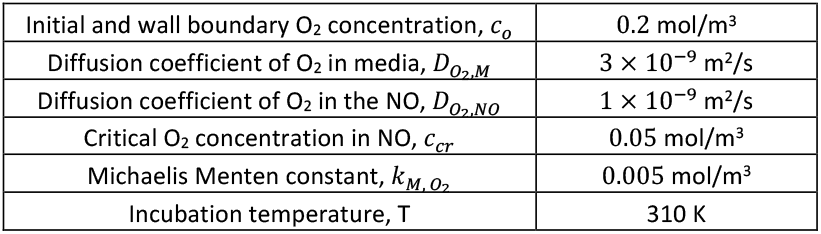
Model parameters for diffusion-reaction kinetics.

|  |  |
| --- | --- |
| Initial and wall boundary O<sub>2</sub> concentration, $c_o$ | 0.2 mol/m<sup>3</sup> |
| Diffusion coefficient of O<sub>2</sub> in media, $D_{O_2,M}$ | $3 \times 10^{-9}$ m<sup>2</sup>/s |
| Diffusion coefficient of O<sub>2</sub> in the NO, $D_{O_2,NO}$ | $1 \times 10^{-9}$ m<sup>2</sup>/s |
| Critical O<sub>2</sub> concentration in NO, $c_{cr}$ | 0.05 mol/m<sup>3</sup> |
| Michaelis Menten constant, $k_{M,O_2}$ | 0.005 mol/m<sup>3</sup> |
| Incubation temperature, $T$ | 310 K |

We calibrated our model using experimental data. We defined the maximum O_2_ consumption rate *k*_*max*_ using the dimensionless parameter Damköhler number. *Da* is a dimensionless number used to compare the characteristic timescales of reaction kinetics to the timescales of transport processes such as mass transfer ^40,41^. Since the reaction happens only in cells within the NO, we defined *Da* just for the NO domain, as shown below. Since, 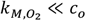, we assume zeroth order kinetics to define *Da*.

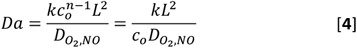

Here, *n* is the order of reaction, *k* is the reaction rate (mol/m^3^s), *c*_*·*_ is the initial O_2_ concentration [mol/m^3^], 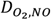 is the diffusion coefficient of O_2_ in the NO [m^2^/s], and *L* is the characteristic diffusion length [m]. Based on the typical values in the literature, we assume *L* to be 300 μm. ^10,11,42,43^ For zeroth order kinetics, *k*≅*k*_*max*_. So, solving for *k*_*max*_

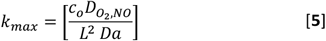

Substituting equation [5] in [3], we modified the Michaelis-Menten kinetic equation.

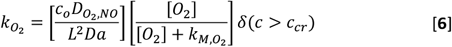

### Modifications to the Damköhler’s number model

To account for the physiologically relevant diffusion limit in our NO, we modified the Damköhler’s model to include the transitions in a diffusion-dominated regime and a reaction-dominated regime.^44,45^ From initial runs to understand consumption in the diffusion-dominated regime, we observed that Damköhler’s number below 0.01 would not affect the O_2_ consumption pattern, as shown in **Figure S1**. Hence, we assumed *Da*_*D*_ = 0.01 as the *Da* along the diffusion-dominated NO surface. We used a ramp function from the NO surface to the radial distance of 300 μm (diffusion limit), bringing up the *Da* to experimentally calibrated *Da*_*R*_ (1-20). We held it constant beyond that. The *Da* of the regimes is defined using a ramp function in COMSOL as:

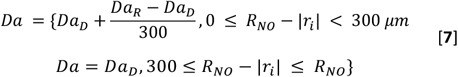

where *R*_*NO*_ is the radius of the NO (μm), *Da*_*D*_ is the *Da*s number on the organoid surface, and *Das* is the *Da* number for the reaction-dominated regime. |*r*_*i*_| is the spatial displacement of the finite element *i*, from the center of the organoid (μm) with (*x*,*y, z*) as coordinates calculated as,

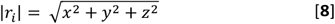

Beyond the diffusion limit, where reaction dominates, we used a constant *Da*_R_ until we reached the center of the NO. We calculated the necrotic area generated at the different *Da*_R_ values to calibrate our model with experimental data.

### Convective transport of media

We coupled our diffusion-reaction model from the static state model with the Laminar flow module in COMSOL to simulate the convective transport of oxygen in the media during dynamic culture using orbital shaking and microfluidic flow around and within the NO. Due to the high cellular density of the NO, we assumed convective transport of O_2_ within it to be negligible, and primarily occurring via diffusion. ^10–13,14^

*For orbital shaking*: To simulate the shaking culture of a NO in shaking culture, we generated a moving mesh to simulate media on an orbital shaker. This mesh represented the media moving around the organoid with a predefined orbital movement. The orbital shaker algorithm updates the coordinates of each finite element to describe the deformation in the media over time, considering the predefined angular velocity with respect to the center of the orbital shaker.

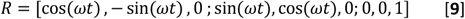

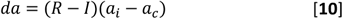

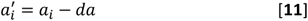

Here, *R* is the rotation matrix, *I* is an identity matrix, *ω* is the angular velocity of the element [rad/s], *a*_*i*_ is the coordinate of the finite element, and *a*_*c*_ is the coordinate of the rotation center. *da* here is the coordinate change, which is used for calculating the updated coordinate of the medium in a rotating frame. For our simulation, we assumed *a*_*c*_ to be [0,0,0], that the orbital shaker has a radius of gyration of 19 mm, just assuming the local movement within a shaking well (six-well plate). Also, 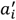 represents the updated coordinate of the finite element. For each deformation of the mesh with respect to time, the coordinate of the element is updated with *da*.

#### For strategies involving microfluidic flow

To simulate flow, we developed a Multiphysics model that coupled the original diffusion-reaction model with a fluid dynamics model. We used the Navier-Stokes equation and assumed the media around the organoid to be an incompressible Newtonian fluid. The Navier-Stokes equations, which describe the motion of fluids like liquids and gases, are given by the continuity equation, which ensures the conservation of mass and momentum to determine the velocity field *u* resulting from the convective transport of the fluid.

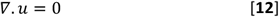

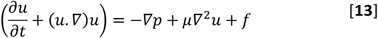

Here, *ρ* is the fluid density [kg/m^3^], *p* is the pressure field [Pa, N/m^2^, kg/m.s^2^], is *μ* the dynamic viscosity [kg/m.s, Pa.s] and *f* represents body or volume forces such as gravity or other external forces [N/m^3^, kg/m^2^.s^2^], which we assume as 0. We list the parameters for convective transport models in **Table 2**.

**Table 2:** Model parameters for diffusion-reaction kinetics.

|  |  |
| --- | --- |
| Initial and wall boundary $O_2$ concentration, $c_o$ | 0.2 mol/$\text{m}^3$ |
| Diffusion coefficient of $O_2$ in media, $D_{O_2, M}$ | $3 \times 10^{-9} \text{ m}^2/\text{s}$ |

### Geometry and boundary conditions

We used 3D models of the NO culture systems housed within a 6-well plate imported from CAD software like Solidworks (Dassault Systèmes). We assumed the NO to be spherical and varied its diameter between 500 and 1000 μm to calculate the necrotic area for each culture strategy. We discretized the finite element model of the spherical NO and surrounding media using an axisymmetric approach with free triangular elements. For the final model, element sizes in the mesh ranged from 0.9 to 47 µm for the NO. We refined the mesh density near the interface between the NO and media to account for steep diffusion gradients, with the finest mesh applied within the NO and a coarser mesh in the media domain. We set the side walls of the media to a fixed constant O_2_ concentration of 0.2 mol/m^3^, and continuity was assumed at the interface between the NO and media. We assumed a diffusion length of 300 µm. We modeled O_2_ uptake to occur exclusively within the NO. Additionally, we applied no-flux boundary conditions to the bottom and top boundaries of the well plate. The initial O_2_ concentration was set uniformly at 0.2 mol/m^3^ throughout the domain. The details of the mesh convergence study, and element sizing are detailed in **Figures S3 and S4** and **Table S1**.

## Results and discussion

### Calibration of Damköhler’s number with experimental data

To evaluate the effects of mechanical culture conditions on viability, we experimentally cultured NOs under two distinct conditions: static culture and orbital shaking (88 RPM). We tested each culture condition using a sample set of n = 3, with NOs reaching an average diameter of *≈* 650 μm. We observed that the NOs maintained under long-term static culture conditions exhibited substantial necrosis, occupying an average of 24% of the total NO area. In contrast, NOs cultured under orbital shaking at 88 RPM displayed significantly higher viability, with only ∼1% necrotic area observed across samples. (**Figure 2A, B)** Additional details about the immunohistochemistry conducted are in the supplementary information sections **S3-S6 and Figure S10**.

**Figure 2.**
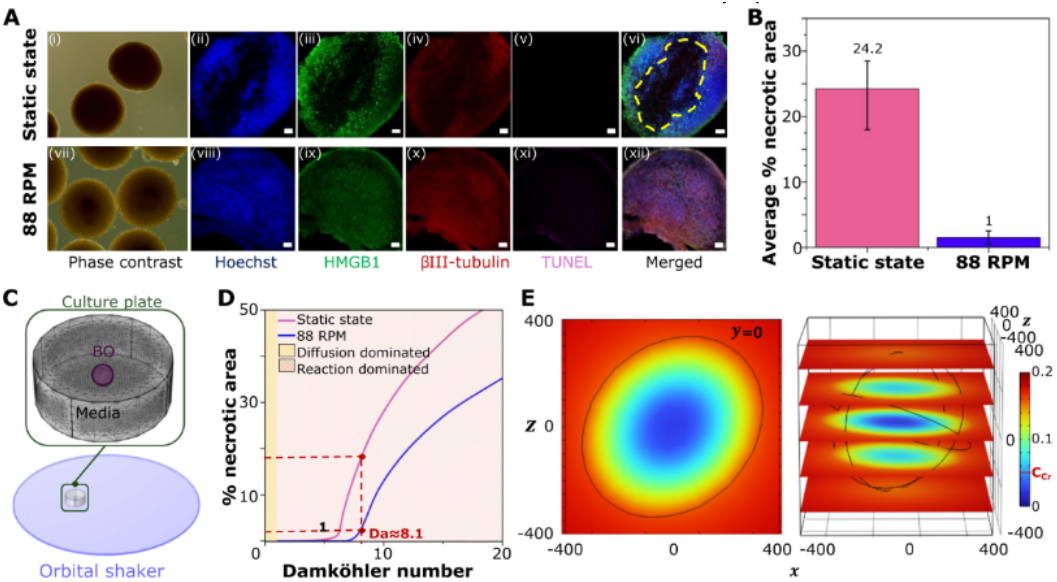
Calibration of the model using experimental NO culture data. (**A**) Confocal microscopy of NO cultures at (**i-vi**) static state, (**vii-xii**) oOrbital shaking at 88 RPM. Nuclei stained with Hoechst 33342 (*blue*), neuronal staining with HMGB-1 (*green*) and βIII-tubulin (*red*), and apoptosis markers with TUNEL (*pink*). A yellow dotted line indicates the necrotic area. The scale bar is 50 μm. (**B**) Experimental estimation of percentage (%) necrotic area of NO. The sample size used was n=3. (**C**) Geometry and mesh of the calibration model; (**D**) Calibration curve of % percentage necrotic area vs Da across diffusion and reaction dominated regions. Using the % necrotic area of NO at static state and 88 RPM as shown in (**B**), we find a corresponding *Da*. We found reasonable agreement with both experimental ranges at *Da≈* 8.1. (**E**) 3D simulation showing oxygen consumption and necrosis across an irregular NO (like experimental sample) with equivalent radius (650 μm) at *Da*= 8.1. Critical oxygen concentration is assumed to be 0.05 mol/m^3^.

We calibrated our model to the experiment by building a 3D culture geometry using the size specifications of a culture well in a six-well plate filled with media (with a growth area of 9.6 cm^2^ and media volume of 2 mL, as shown in **Figure 2C**. We assumed the initial O_2_ concentration in the NO and culture plate as 0.2 mol/m^3^ throughout the domain. We built a spherical NO matching the average of previously cultured NOs (diameter = 650 μm, cultured in a well of a six-well plate with a gyration radius of the orbital shaker of 19 mm). We ran a parameter sweep on our COMSOL model to run at various Damköhler numbers (*Da*_R_ = 0.1-20). We plotted the percentage necrotic area observed to generate a calibration curve for (88 RPM) and static state (0 RPM), as shown in **Figure 2D**. We ran all our culture simulations using the same well size (six-well plate) and shaking configuration (19 mm radius of gyration) to correlate with the experimental conditions.

To get a good agreement of the *Da* across both static and 88 RPM conditions, we aligned the lower range of necrosis observed experimentally in static cultures with the average necrosis value in shaking cultures. This yielded a reaction regime *Da* of *Da*_R_ = 8.1 for our model. (**Figure S2**) To validate our calibrated model, we simulated the 3D structure of an irregular NO under static conditions based on **Figure 2A (vi)**, which produced 18.2% necrosis compared to the average of 21% necrosis observed in the experimental data. The simulated necrotic regions closely resembled experimental findings in shape and size, as in our NO slice.

We further validated the robustness and adaptability of our model by extending it to previously published regional-specific neural organoid cultures exposed to a range of orbital shaking conditions (0–110 RPM). Using our generated calibration curves (**Figure S8**), we accurately reproduced the shape, location, and extent of necrotic regions reported in the literature, even in geometrically irregular NOs. The strong agreement between simulated and experimental outcomes confirmed the model’s predictive capability not only for assessing oxygen diffusion limits and necrosis severity but also for locating and spatially mapping necrotic zones. These results highlight the utility of our framework as a powerful tool for benchmarking and optimizing various NO culture strategies.

### Conventional NO culture methods are limited in reducing necrosis

Each conventional culture strategy used for NOs relied on diffusion-dominated or forced convection-enhanced transport. Hence, it was crucial to determine how these mechanisms influence O_2_ and nutrient availability at different NO sizes. In static cultures, transport was purely diffusion-driven, governed by Fick’s laws for O_2_. In shaking conditions, fluid motion enhances mass transport by reducing the resistance to diffusion and improving O_2_ and nutrient exchange. Microfluidic flow, in contrast, introduced controlled convective transport, where flow velocity profiles dictated shear forces and nutrient gradients. By systematically assessing the progression of necrosis as the NO grows using these models, we aimed to establish the upper size limits for maintaining viable NOs under each culture strategy. (**Figure 3A-C**). We ran simulations for three NO sizes-500, 650, and 1000 μm diameter. We note that in all cases, the modeled overall volume of media in the well (standard six-well plate) was 2 mL.

**Figure 3.**
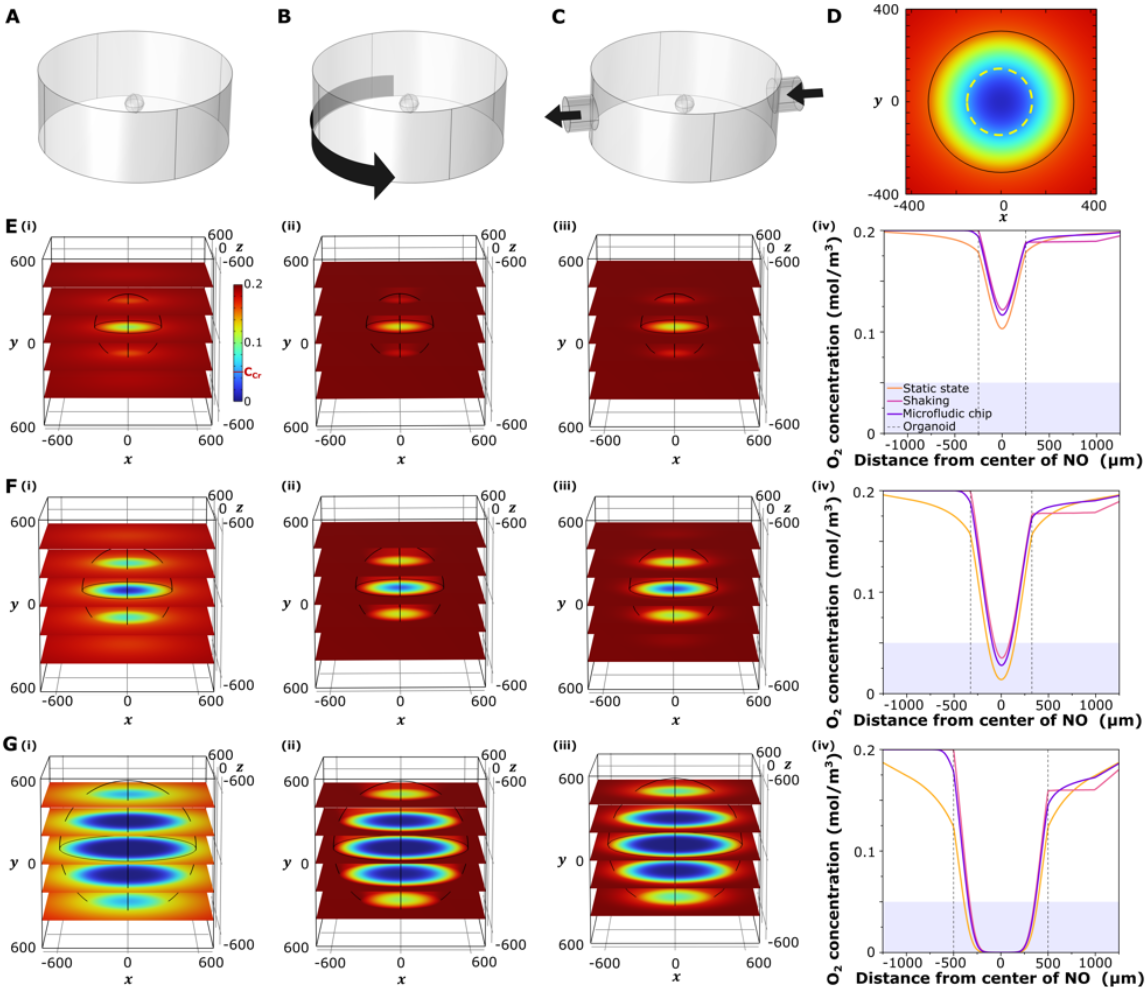
Comparison of necrosis across different NO culture strategies. **(A-C)** 3D geometry of NO culture strategy using **(A)** static culture, **(B)** orbital shaking, and **(C)** microfluidic flow around the NO. **(D)** Visualization of the necrotic area by plotting O_2_ concentration in a NO slice. The core area marked by the dotted yellow line has an O_2_ concentration below the critical O_2_ concentration of 0.05 mol/m^3^ and is necrotic. **(D-F)** Computational estimation of necrosis in a NO by plotting O_2_ concentration across an NO between 500-1000 *μ*m (diameter). (**D**) For a 500 *μ*m NO cultured using, **(i)** static culture **(ii)** orbital shaking, and **(iii)** microfluidic flow around NO. **(iv)** O_2_ concentration across NO on a cutline at y=0 for all three cases. The blue-shaded region represents the area below the critical O_2_ concentration, which determines the necrotic area. We observe viable NOs with no necrosis in all culture strategies. **(E)** For a 650 *μ*m NO cultured using **(i)** static culture **(ii)** orbital shaking, and **(iii)** microfluidic flow around NO. **(iv)** O_2_ concentration across NO on a cutline at y=0 for all three cases. The blue-shaded region represents the area below the critical O_2_ concentration, which determines the necrotic area. We observe that orbital shaking improves the NO viability in comparison to the other two culture strategies. **(F)** For a 1000 *μ*m NO cultured using, **(i)** static culture, **(ii)** orbital shaking, and **(iii)** microfluidic flow around NO. **(iv)** O_2_ concentration across the NO on a cutline at y=0 for all three cases. The blue-shaded region represents the area below the critical O_2_ concentration, which determines the necrotic area. We observe that all strategies show significant necrotic cores.

We optimized our mesh for each strategy to ensure consistent results while minimizing computational time. Specifically, we identified the coarsest mesh that produced results identical to those obtained with a finer mesh. To achieve this, we optimized two parameters: the number of layers at the interface between the medium and the NO and the overall predefined physics-controlled mesh size for the NO (**Figure S3**). After running mesh convergence tests for each culture condition, we observed an 8-layer boundary mesh combined with an extra fine predefined mesh for static cultures and an 8-layer boundary mesh with a finer predefined mesh for shaking and microfluidic cultures, which yielded stable results with minimal computational cost (**Figure S4, 5**). We maintained the shaking speed at 88 RPM, and for microfluidic flow, we assumed an initial flow rate of 200 μL/min, as seen in conventional literature. ^25^

We observed that NOs 500 μm in diameter remained stable across all culture strategies, suggesting that O_2_ and nutrient diffusion were sufficient to sustain viability. This aligned with theoretical diffusion limits, as the radially symmetric structure of spherical NOs allows for relatively uniform nutrient distribution within this size range. When the NO size increased to 650 μm, necrosis emerged across all three culture strategies, though to varying degrees. Shaking culture exhibited the least necrosis at *≈*5%, while microfluidic flow resulted in *≈*9% necrosis, and static culture showed the highest necrosis at *≈*19%. These findings indicated that forced convection extended the viability of NOs beyond passive diffusion limits but did not entirely prevent necrotic core formation. At 1000 μm, simulations revealed significant necrosis (>45%) in all culture strategies, highlighting a fundamental size limitation regardless of the transport method. This suggested that while shaking culture improved NO viability, none of the conventional culture strategies effectively sustained NOs beyond 650-700 μm diameter due to inherent diffusion constraints.

More detailed plots illustrating necrosis progression with increasing NO size under each condition are provided in **Figure S5**. We also studied the effect of flow rate in microfluidic flow around NOs (650, 1000 μm diameter). We observed that while microfluidic flow reduced necrosis significantly in the case of 650 μm (18% to 9%), the necrosis remained significant, between 50-60%, in the case of a 1000 μm NO. (**Figure S6)**

### 3D spatial perfusion within NO eliminates necrosis

While conventional culture methods resulted in necrosis beyond a critical organoid size, we hypothesized that introducing internal spatial perfusion within the NO could mitigate and potentially prevent necrosis across NOs of varying size scales. To test this hypothesis, we developed computational models incorporating embedded 3D synthetic microfluidic channels or capillaries within NOs to facilitate synthetic vasculature-like perfusion. We modeled the capillaries with gel-like properties using the gelatin built-in module in COMSOL to simulate physiologically relevant diffusion and mechanical characteristics, as shown in **Table 3**. To account for potential bulk media accumulation around the NO, we assumed that the concentration at the interface between the NO surface and the surrounding bulk media (wall) remained constant at 0.2 mol/m^3^. Furthermore, to better reflect the influence of internal perfusion, we redefined the *Da* variation relative to the capillary surface because it served as the direct source of fresh media to cells. This contrasts with our previous models, where *Da* was defined relative to the surface of the NO, assuming a diffusion-dominated regime followed by a reaction-dominated regime. (**Figure S7**).

**Table 3:** Model parameters for internal spatial perfusion of NO.

|  |  |
| --- | --- |
| Diffusion coefficient of $\text{O}_2$ in the capillary wall | $2 \cdot 10^{-9} \text{ m}^2/\text{s}$ |
| Capillary diameter | 20 $\mu\text{m}$ |
| Capillary wall thickness | 5 $\mu\text{m}$ |
| Capillary spacing | 350-550 $\mu\text{m}$ |
| Equivalent diameter of NO | 1200 $\mu\text{m}$ |

The first parameter we optimized was the capillary distance to ensure adequate O_2_ and nutrient delivery within NO to meet the consumption demands. We simplified the computational model, by designing a two-capillary geometry embedded within a cubical organoid and systematically varied the capillary distance between 350 and 550 μm to analyze diffusion-reaction trends (**Figure S7**). Despite the theoretical diffusion length of 300 μm, our simulations indicated that meeting cellular uptake demands required a significant overlap between the diffusion regimes of adjacent capillaries to maintain a stable concentration gradient. Through this analysis, for our NO system, we determined that the optimal capillary distance— measured from capillary surface to surface—should be approximately 400 μm, ensuring efficient diffusion while minimizing necrotic regions.

Building on our findings, we simulated a 1200 μm NO with spatial perfusion varying the capillary density between one capillary, four capillaries, and nine capillaries to evaluate their effectiveness in reducing/preventing necrosis (**Figure 4**). The one-capillary (49%) and four-capillary systems (33%) reduced necrosis compared to the static state simulation (60%). Still, they failed to meet the NO’s metabolic demands, leading to large necrotic regions. This insufficiency was primarily due to the limited overlap of diffusion regions between the capillaries, resulting in O_2_-deprived zones between them.

**Figure 4.**
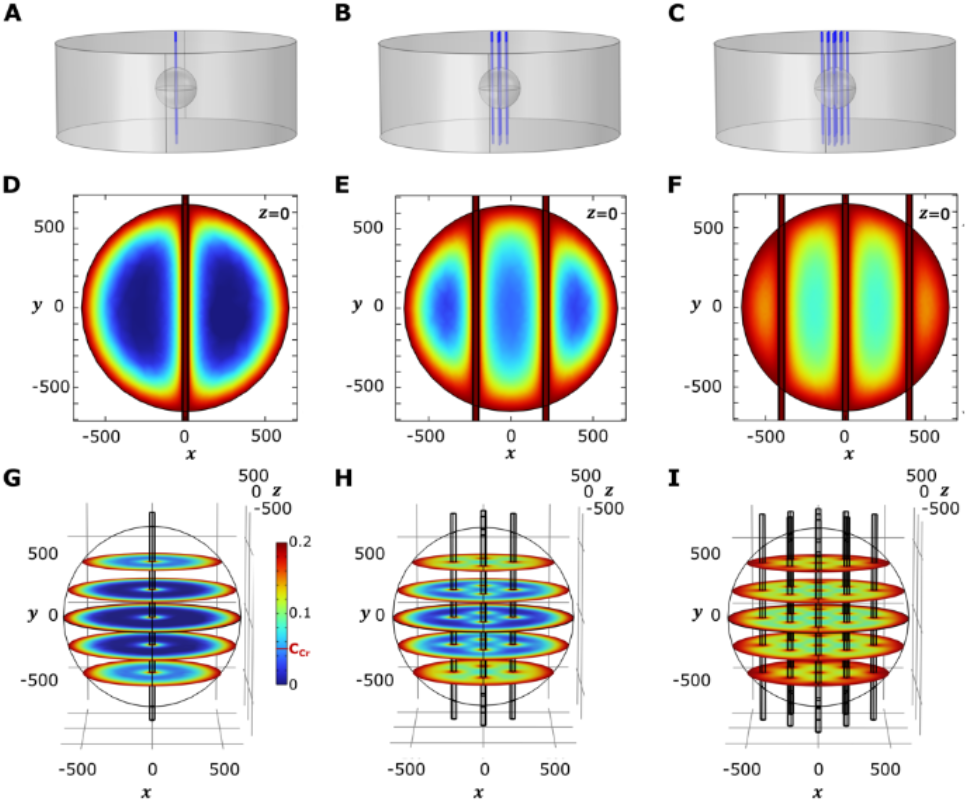
Microfluidic perfusion within NO prevents necrosis in a NO with an equivalent diameter ∼1200 μm. **(A-C)** Varying synthetic capillary density to determine optimal layout to eliminate necrosis. We varied the density in the grid layout containing **(A)** 1, **(B)** 4, and **(C)** 9 capillaries. We maintain a flow rate of 200 μL/min. **(D-F)** O_2_ concentration across NO on a cutline at z=0 for the three capillary densities. **(G-I)** Stacked 3D visual representation of O_2_ concentration across the NO for the three capillary densities. **(D&G)** 1 capillary system causes high necrosis. (**E&H**) 4 capillary system necrosis reduces, but the organoid is not viable. **(F&I)** We observe no necrosis in the 9-capillary system.

In contrast, the nine-capillary system effectively maintained O_2_ availability throughout the NO. The lowest O_2_ concentration in the NO remained just above the critical threshold (0.05 mol/m^3^), ensuring a fully viable non-necrotic NO. These results demonstrated that a higher density of internal perfusion capillaries significantly improved O_2_ distribution, making large-scale NO cultures feasible without necrosis.

We further demonstrated the robustness and scalability of our model and design strategy by conducting preliminary simulations using alternative piping layouts. Specifically, we modeled a 1000 μm NO with a branched, vasculature-inspired channel architecture. By varying the capillary spacing within the range from 240 μm to 300 μm, we achieved uniform perfusion throughout the organoid, with no necrotic regions observed (Lowest O_2_ concentration: 0.121 mol/m^3^). (**Figure S9A**) Carrying this idea forward, and to demonstrate scalability, we also designed an alternate channel layout that was inspired by a tree-like branching gradient to support a 2600 μm NO with capillary spacing from 60 μm to 530 μm. The design had two inlets, one with just media and another with media mixed with an arbitrary biochemical input. The tree-like piping network uniformly perfused the organoid, while the dual-inlet configuration established a biochemical gradient. This simulation demonstrates the potential of our strategy to simultaneously enable media perfusion and spatial biochemical patterning, supporting the generation of large, bio-complex, and viable NOs (Lowest O_2_ concentration: 0.085 mol/m^3^). (**Figure S9B**)

These results supported the ability of our approach to test and scale complex bioreactor designs for larger scale organoid cultures.

## Conclusion

We developed a comprehensive 3D finite element model to quantify necrosis in NOs, incorporating convection, diffusion, and reaction dynamics across conventional culture techniques, including static, orbital shaking, and microfluidic flow around NO. Our computationally intensive approach captured necrosis in 3D CAD-based geometries, providing a more accurate representation of the 3D tissue environment in conventional culture strategies. We accurately captured the orbital shaking culture strategy by implementing a moving mesh approach, ensuring precise simulation of fluid dynamics and convective transport. Additionally, rigorous mesh convergence testing enabled optimal mesh sizing, balancing accuracy with computational efficiency. We developed a new calibration method by utilizing the Damköhler number to adjust the reaction-to-diffusion dependency in our (NO) model. We enhanced the applicability of the model to diverse cell culture systems by generating calibration curves for commonly used methods, including static culture and orbital shaking across a speed range of 2–110 RPM. Applying this calibration approach, we achieved a reasonable agreement between our model predictions and experimental NO cultures at static conditions and 88 RPM orbital shaking, demonstrating the robustness and adaptability of our method.

Our model demonstrated that while orbital shaking reduces necrosis more effectively than static culture and microfluidic flow, it remains insufficient for sustaining NOs larger than 700 μm in diameter. We suggest a strategy to overcome this limitation using a device design incorporating embedded synthetic capillaries within NOs, allowing them to grow around them while being perfused. We optimized the capillary spacing to be around 400 μm. This approach aims to enhance oxygen and nutrient delivery, mitigate necrosis in larger NOs, and improve long-term viability. We also demonstrated preliminary simulations of chemical patterning and alternate vasculature-like layouts.

In the future, we will expand our framework to calibrate the model and optimize design parameters, such as capillary geometry, layout, and density, to match the metabolic demands of specific cell cultures. Additionally, we plan to extend the model to account for other critical metabolites beyond oxygen, further enhancing its applicability to diverse biological systems. This work opens up new design possibilities for 3D perfusive microbioreactors, which can be fabricated using high resolution 3D fabrication technologies (two photon polymerization, 3D printing etc) to enable more sophisticated architectures, including parallel, serpentine, and helical channel configurations. These easily scalable bioreactors would offer not only improved reductions in necrosis but also the ability to guide spatiotemporal tissue differentiation through the generation of precise local and global biochemical gradients. We envision that this modeling framework will extend well beyond NOs, serving as a versatile platform for designing next-generation bioreactors, 3D tissue engineering systems, organoid-based research, and engineered living materials.

## Author contributions

**AP**: Conceptualization, data curation, formal analysis, investigation, methodology, validation, visualization, writing – original draft, writing – review & editing

**BZ**: Data curation, formal analysis, investigation, methodology, validation, visualization, writing – original draft, writing – review & editing

**IEMP**: Conceptualization, data curation, formal analysis, investigation, methodology, validation, visualization, writing – original draft, writing – review & editing

**AF**: Visualization, original draft, writing – review & editing

**DG**: Conceptualization, writing – review & editing

**DMA:** methodology and investigation

**AL:** methodology and investigation

**LS**: Funding acquisition, supervision, conceptualization, data curation, writing – original draft, writing – review & editing

**DHG**: Funding acquisition, supervision, conceptualization, formal analysis, methodology, validation, visualization, writing – original draft, writing – review & editing.

## Conflicts of interest

LS is a consultant for AxoSim, New Orleans.

## Data availability

The data supporting this article have been included as part of the Supplementary Information. Additional data are available upon request.

## Acknowledgements

Research reported in this publication was supported by the Johns Hopkins University SURPASS Program and NSF (EFMA 2318093). IEMP was supported by NIH (K12-GM123914) and the Colgate-Palmolive Award in In Vitro Toxicology. DMA was supported by the NIH (T32 ES007141) and the International Foundation for Ethical Research Graduate Fellowship.

## Notes

### Competing Interest Statement

LS is consultant for AxoSim, New Orleans

