## Supplementary Information for "Computational modeling of necrosis in neural organoids"

#### S1 Model setup, mesh configuration, and simulation data

##### S1.1 Diffusion regime calibration

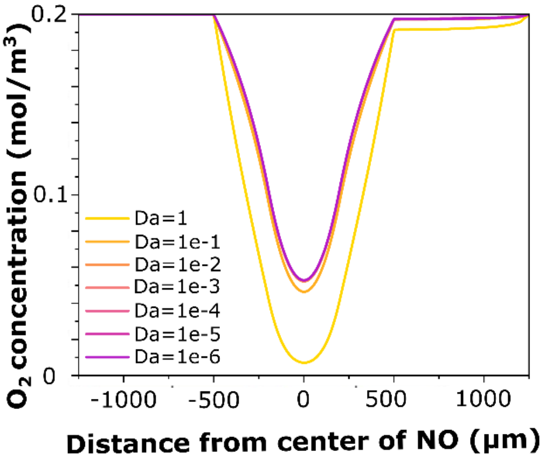


**Figure S1 Variation of diffusion regime Damköhler (*Da*) numbers.** $O_2$ concentration was plotted across a cutline at *y*=0 the in a 1000 μm NO.

We determined the boundary condition Damköhler number ($Da_D$) for diffusion-only transport at the edge of the NO, by simulating a 1000 μm NO for various *Da* in the diffusion-dominated region ($1e^{-6}$ to 1) to find a stable value of Da where the reaction is negligible. We observed convergence issues resulting from a sudden increase from *Da*=0, which resulted in numerical instabilities. We also observed that for Da values below 0.01, variations in Da had a negligible effect on the oxygen consumption profiles within the NO. To avoid any numerical instabilities and convergence issues, we chose to use the maximum possible value $Da_D$ = 0.01 to establish an initial ramping condition.

##### S1.2 Necrotic area estimation

We chose five cut planes for each NO, from the center of the NO to 400 μm and 800 μm for visualization of the necrosis in the NOs. We kept the frames all the same for all NOs, regardless of size. Using the center cut plane, we assumed symmetry and extracted the $x, y$, and $c$ data from the simulation across the cut line within the NO at ($z = 0$) to generate our $O_2$ gradient curves. Here $x, y, z$ are linear coordinates, and $c$ is the $O_2$ concentration. To calculate the necrotic area in the NO, we identified and counted the coordinates where ($c < c_{cr}$) as the number of necrotic or necrotizing regions $m$. Since the element sizes vary minimally in the NO, we assumed them to be uniform to calculate the unit lengths $dx$ and $dy$ given as:

$$dx = \frac{total\ range\ of\ x\ values}{number\ of\ x\ values} \quad [S1]$$

$$dy = \frac{total\ range\ of\ y\ values}{number\ of\ y\ values} \quad [S2]$$

We calculated the unit area per necrotic region as:

$$unit\ area = dx \times dy \quad [S3]$$

Finally, we calculated the necrotic area $A_n$ as:

$$A_n = n_o \times unit\ area \quad [S4]$$

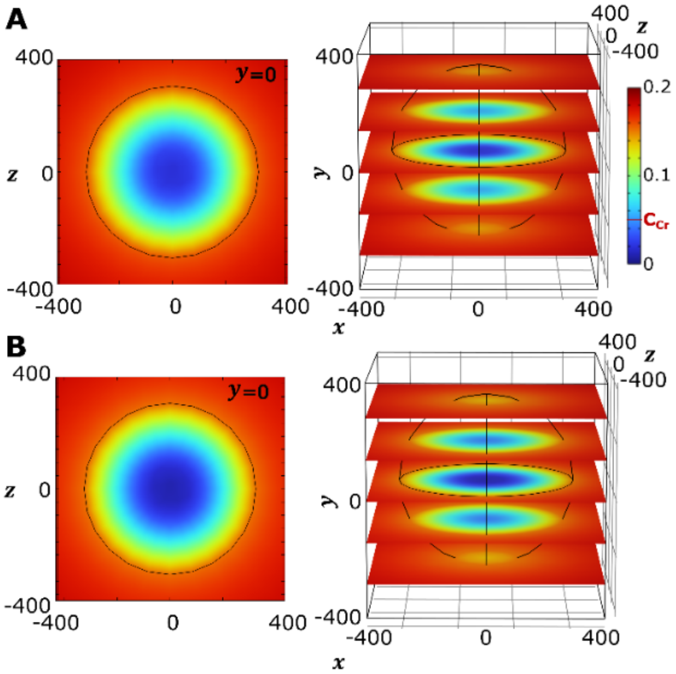


**Figure S2 Range of necrosis in an NO cultured in static state conditions.** (**A**) NO with 18% necrosis at Da=8 and (**B**) 24% necrosis at Da=9.1. We chose a Da value close to (**A**) to get a reasonable agreement between necrosis observed in experimental NOs cultured using static and orbital shaking in the final model.

##### S1.3 Mesh convergence studies

We used a tetrahedral mesh and varied the element sizing within the NO (diameter 650 μm) using the various mesh sizes in COMSOL. Furthermore, we adjusted the mesh element size by implementing between 2 and 8 boundary layers, allowing for a detailed investigation of mesh resolution effects. (**Figure S3**) Mesh sizing details are mentioned in **Table S1**.

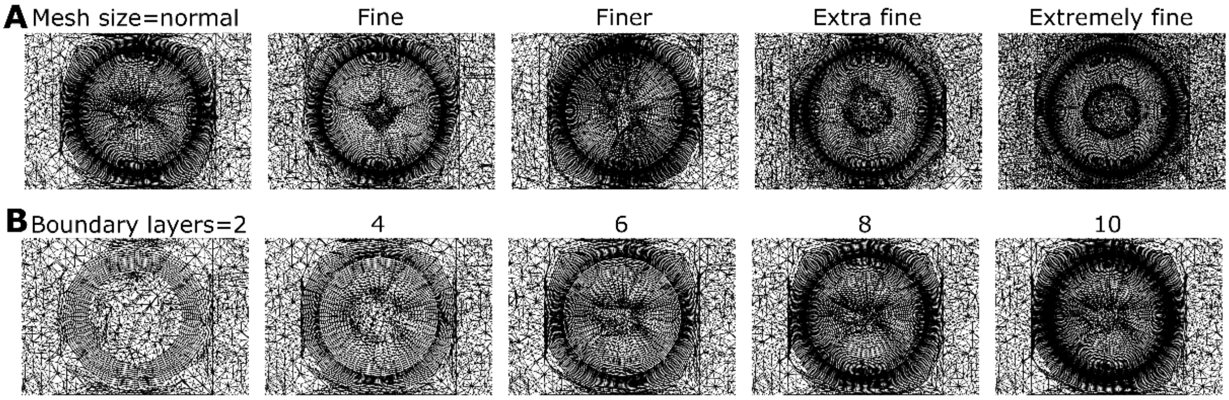


**Figure S3 Mesh parameter variation by** (**A**) varying size of NO mesh elements and (**B**) varying number of boundary layers on the NO-media interface.

**Table S1: Element sizing in each mesh configuration within NO**

| **Mesh** | **Maximum element size (μm)** | **Minimum element size (μm)** | **Maximum element growth rate** | **Curvature factor** | **Resolution of narrow regions** |
|---|---|---|---|---|---|
| Normal | 85 | 8 | 1.5 | 0.6 | 0.5 |
| Fine | 80 | 7 | 1.45 | 0.5 | 0.6 |
| Finer | 63 | 4.5 | 1.4 | 0.4 | 0.7 |
| Extra fine | 47 | 3 | 1.35 | 0.3 | 0.85 |
| Extremely fine | 23 | 0.9 | 1.3 | 0.2 | 1 |

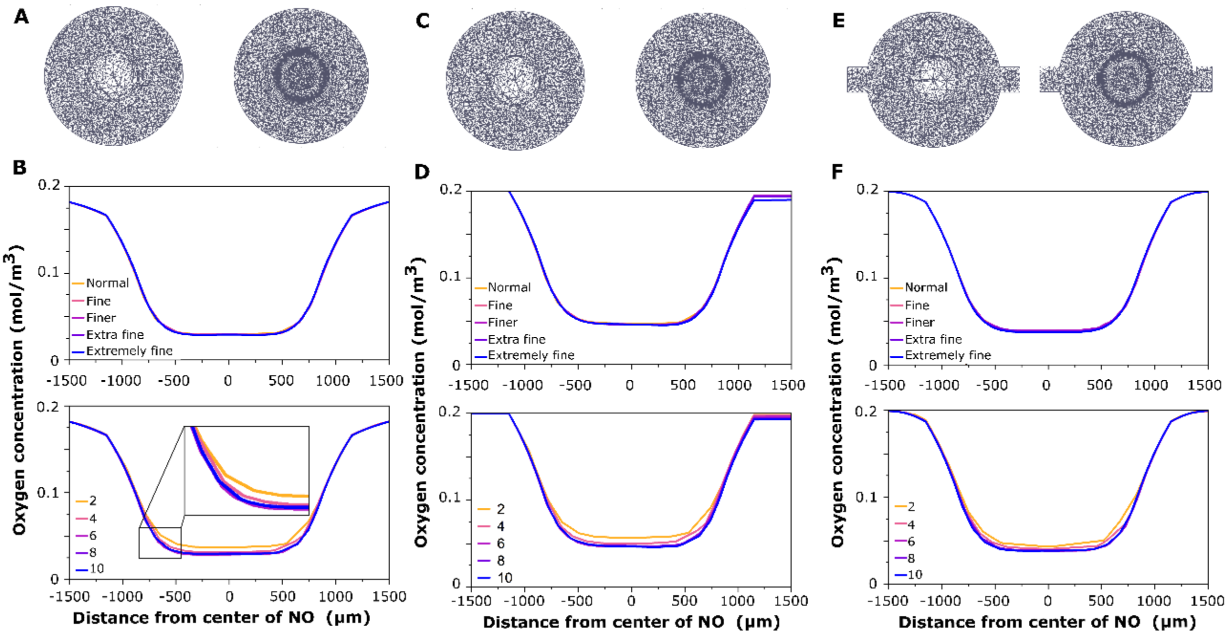


**Figure S4 Mesh convergence study for NOs cultured using different conditions while varying mesh sizes and boundary layers**. (**A-B**) Culture using static state culture. (**A**)The optimal mesh for achieving consistent oxygen consumption profiles while minimizing computational time. (**B**) $O_2$ concentration across the NO on a cutline at y=0. (**C-D**) Culture using orbital shaking. (**C**) The optimal mesh for achieving consistent oxygen consumption profiles while minimizing computational time. (**D**) $O_2$ concentration across the NO at y=0. (**E-F**) Culture using microfluidic perfusion around the NO. (**E**) Optimal mesh layout for microfluidic perfusion around the NO. (**F**) $O_2$ concentration across the NO on a cutline at y=0.

We obtained $O_2$ consumption profiles in different culture strategies with the meshes defined in **Figure S3**. We kept the mesh size in the surrounding media at a normal size setting to reduce computational time, as the media itself was not the primary focus. For orbital shaking conditions, we considered a well in a six-well plate on an orbital shaker with a gyratory radius of 19mm. Instead, we applied boundary layer refinements at the NO-media interface to ensure adequate resolution. In static culture, we employed an extra-fine predefined organoid mesh size along with eight boundary layer refinements, which yielded stable $O_2$ consumption profiles. Likewise, we also observed that a finer predefined organoid mesh size, combined with eight boundary layer refinements, maintained consistency in orbital shaking and microfluidic flow around NO. Based on these findings, we selected an extra-fine organoid mesh size and eight boundary layer refinements for all subsequent simulations to ensure reliable and accurate modeling of oxygen consumption.

#### S1.4 Simulations to determine NO necrosis

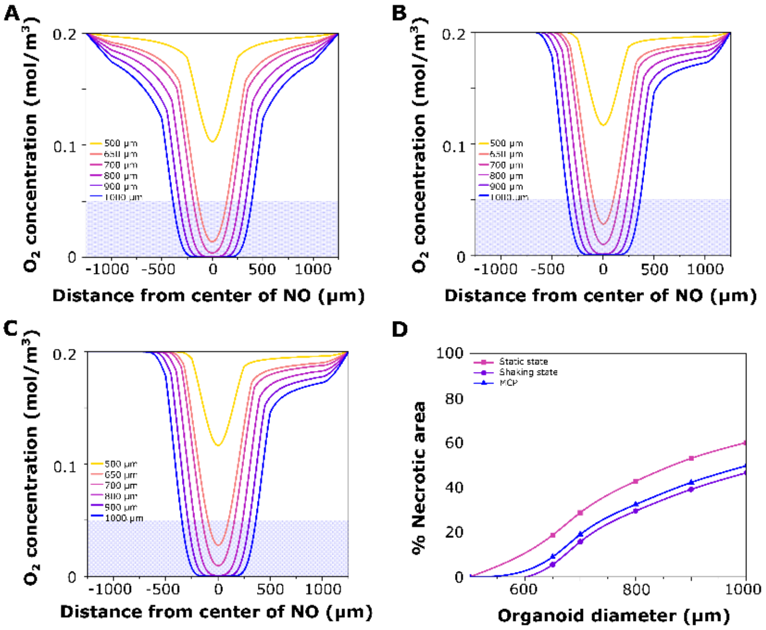


**Figure S5 Necrosis progression as the NO grows.** $O_2$ concentration across NO on a cutline at y=0 cultured using, (**A**) static culture, (**B**) orbital shaking, and (**C**) microfluidic perfusion around the NO. The blue-shaded region represents the area below the critical $O_2$ concentration, which determines the necrotic area. (**D**) % necrotic area for each conventional culture strategy as the NO grows from 500-1000 μm.

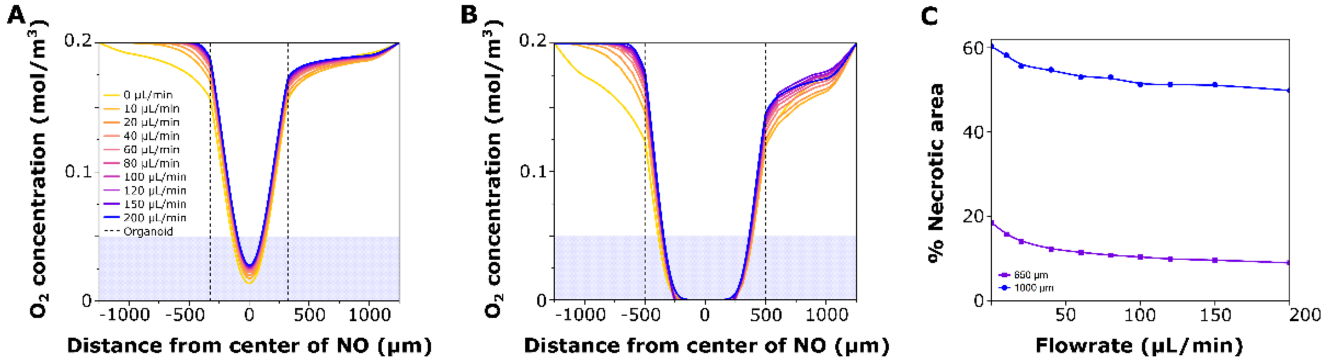


**Figure S6 Necrosis progression with variation in flow rate of microfluidic perfusion of media around NO** of diameter, (**A**) 650 μm, and (**B**) 1000 μm. (**C**) % necrotic area for different flow rates as the NO grows from 650 to 1000 μm.

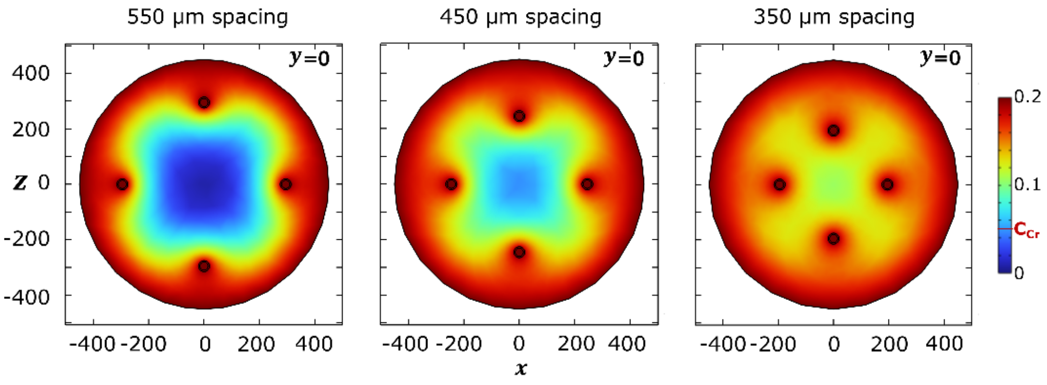


**Figure S7 Necrosis progression with variation in capillary spacing for media flow within NO.** We used a four-capillary grid to feed an NO of diameter 900 μm. We observed significant necrosis of 12.09% with a capillary spacing of 550 μm, minimal necrosis of 0.01% with 450 μm, and a fully viable NO for a capillary spacing of 350 μm.

#### S2 Preliminary studies

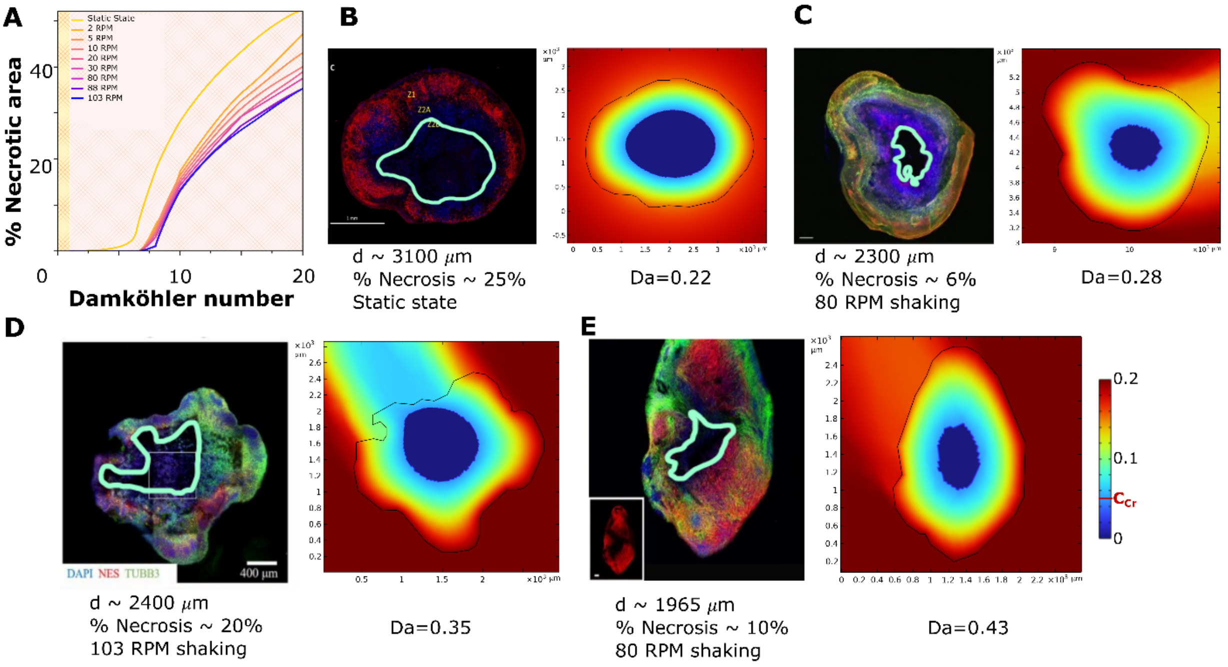


**Figure S8 Calibration and modeling of irregular organoids.** (**A**) Calibration curve of % necrosis vs. Da for static and different orbital shaking speeds relevant to the literature between (2-103 RPM). Using the % necrotic area of NO, we find a corresponding Da that is in agreement with the slice data. (**B**) Microscopy image of an irregular necrotic cerebral organoid. Adapted with permission under the terms of the Creative Commons CC BY license [28] Copyright 2021, The Authors. (*left*), Calibrated simulation using the static state curve in panel (**A**) at Da=0.28 (*right*). (**C**) Microscopy image of an irregular human midbrain organoid with a substantial dead/necrotic core. Adapted with permission under the terms of the Creative Common CC BY license [29] Copyright 2021, The Authors (*left*). Calibrated simulation using the 80 RPM curve in (**A**) at Da=0.28 (*right*). (**D**) Microscopy image of an irregular necrotic brain organoid. Adapted with permission from [32] Copyright 2021, IOP Publishing Ltd (*left*). Calibrated simulation using the 103 RPM curve in (**A**) at Da= 0.45 (*right*). (**E**) Microscopy image of an irregular necrotic cerebral organoid. Adapted with permission from [33] Copyright 2017, Elsevier (*left*). Calibrated simulation using the 80 RPM curve in (**A**) at Da= 0.45 (*right*).

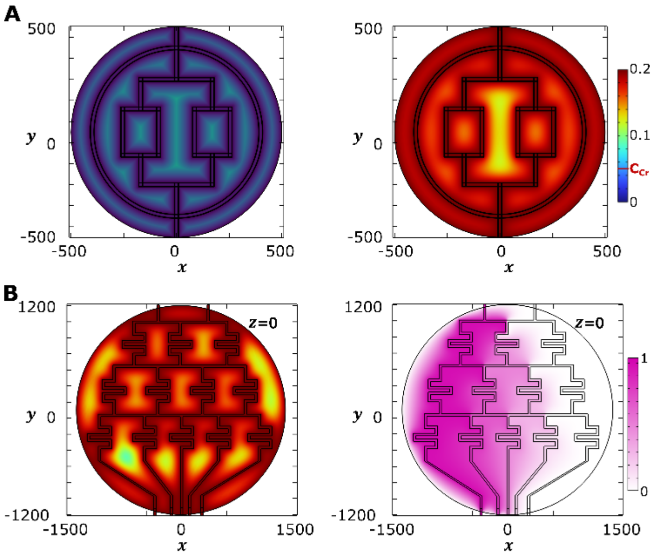


**Figure S9 2D simulations of alternate synthetic capillary layouts for embedded microfluidic flow within NO.** (**A**) Vasculature-like synthetic capillary layout at t=0 (*left*) and at steady state (*right*). (**B**) Chemical gradient generation along with spatial perfusion. $O_2$ concentration across the NO in the device (*left*) and gradient profile (indicated in pink, ranging between 0-1 AU of biochemical, *right*) at steady state.

### S3 NO culture studies

#### S3.1 Immunohistochemistry of NOs

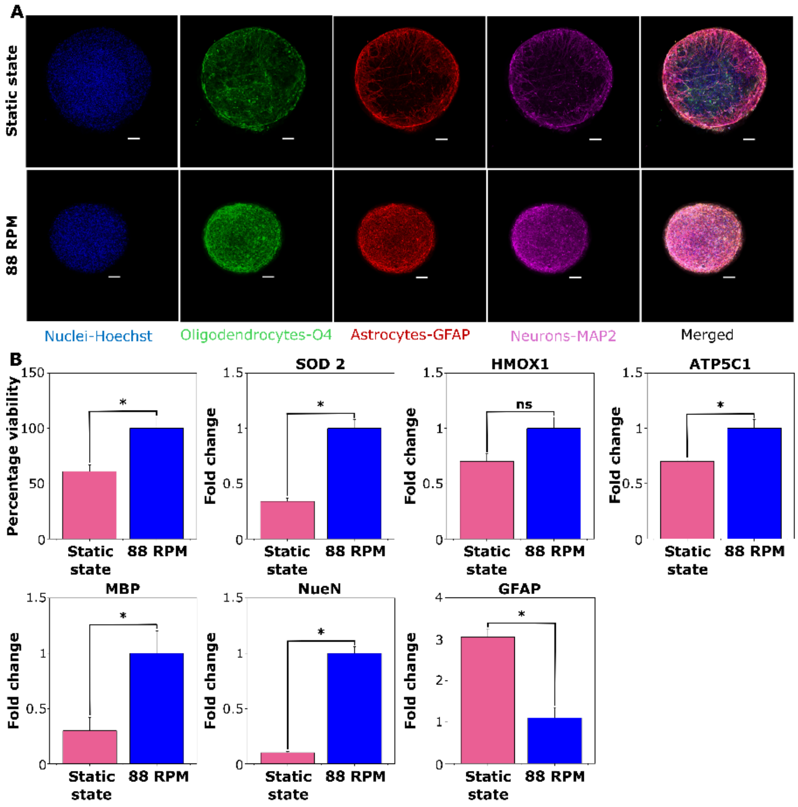


**Figure S10 Cell vulnerability to mechanical stress based on culture conditions: Static culturevs orbital shaking (88 RPM) vs .** (**A**) Representative images of NOs subjected to static conditions (top row) or orbital shaking at 88 RPM and gyratory radius 19mm (bottom row) for immunostaining of various cell types, including nuclei (Hoechst, blue), oligodendrocytes (O4, green), astrocytes (GFAP, red), neurons (MAP2, purple), and the merged image showing all cell types. Scale bars represent 50 μm; (B) Quantification of cell viability and gene expression under static and shaking conditions. Viability, assessed via resazurin assay, was significantly lower in static cultures compared to shaking (median: 64.1% vs. 97.2%; $p = 0.029$, Mann-Whitney U test). Gene expression fold changes for SOD2, HMOX1, ATP5C1, MBP, NeuN, and GFAP are shown. We performed statistical comparisons using the Mann-Whitney U test on four biological replicates per condition (each representing an average of 15 pooled NOs). We observed significant reductions in SOD2, ATP5C1, NeuN, MBP, while upregulating GFAP expression in static cultures ($p < 0.05$). HMOX1 expression showed a downward trend in static conditions but did not reach statistical significance. Data are presented as median ± range, with statistical significance denoted by asterisks ($p < 0.05$).

We cultured NOs under two mechanical conditions: static and orbital shaking at 88 RPM. Gene expression analysis revealed that static conditions significantly reduced the expression of oxidative stress and mitochondrial function genes, including ATP5C1 and SOD2, while HMOX1 trended in the same direction but was not statistically significant. Static culture also significantly downregulated MBP (oligodendrocyte marker) and NeuN (neuronal marker), while upregulating GFAP (astrocytic marker). Immunostaining showed structural abnormalities, such as the necrotic core formation. (**Figure S10, Table S2**).

In follow-up viability assays (**Figure 2A–B**), NOs cultured under static conditions for two weeks exhibited a ~40% reduction in viability compared to shaking controls. TUNEL staining was minimal, indicating low levels of apoptosis in control NOs. HMGB1, a non-histone nuclear protein associated with necrotic and inflammatory signaling, showed higher levels in static cultures. Cells ubiquitously expressed HMGB1, and they released it into the extracellular space during necrotic but not apoptotic cell death. Additionally, glial cells and neurons secrete HMGB1 in response to inflammation such as stroke, traumatic brain injury, and epilepsy [34,46–48]. Although we did not assess HMGB1 localization, elevated expression may reflect cellular stress or necrotic processes. Taken together, these findings suggest that reduced mechanical agitation impairs NO health by disrupting oxidative stress response mechanisms, mitochondrial function, and cell-type-specific marker expression, likely contributing to cell death and loss of structural integrity.

**Table S2**: Primary antibodies used for immunofluorescence

| Antibody or dye | Host | Type | Source / cat no. | Dilution |
|---|---|---|---|---|
| β-Tubulin class III | Mouse | Monoclonal | Sigma T5076 | 1:1500 |
| GFAP | Rabbit | Polyclonal | DAKO Z0332 | 1:400 |
| HMGB1/HMG-1 | Mouse | Monoclonal | R&D Systems MAB1690 | 1:50 |
| MAP2 | Chicken | Polyclonal | Invitrogen PA1-10005 | 1:5000 |
| NF-H | Chicken | Polyclonal | Invitrogen PA1-10002 | 1:1000 |
| O4 | Mouse | Monoclonal | R&D Systems MAB1326 | 1:200 |
| Click-iT TUNEL staining kit | NA | NA | Invitrogen C-10247 | Multicomponent, see manufacturer specifications |

##### S3.2 RT-qPCR

We isolated total RNA via a Quick-RNA™ Microprep Kit (Zymo Research). We determined RNA quantity and purity using NanoDrop™ 2000c Spectrophotometer (Thermo Fisher Scientific). We reverse-transcribed a total of 500 ng of RNA using M-MLV Reverse Transcriptase and Random Hexamer primers (Promega) according to the manufacturer's instructions. We evaluated the expression of genes using the TaqMan gene expression assay (Applied Biosystems) (Table 2). We performed real-time quantitative PCR (RT-qPCR) using a 7500 Fast Real-Time system machine (Applied Biosystems). We normalized gene expression to the expression of the reference gene Actin beta (ACTB), and the primers used are mentioned in **Table S3**.

**Table S3:** Primers used for RT-qPCR analysis

| Assay name | Assay ID | Assay type | Catalog number |
|---|---|---|---|
| ACTB | Hs01060665 | TaqMan® Gene Expression Assay | 4331182 |
| GFAP | Hs00909233 | TaqMan® Gene Expression Assay | 4331182 |
| MBP | Hs00921945 | TaqMan® Gene Expression Assay | 4331182 |
| RBFOX3 (NeuN) | Hs01370653 | TaqMan® Gene Expression Assay | 4331182 |
| SOD2 | Hs00167309 | TaqMan® Gene Expression Assay | 4331182 |
| ATP5C1 | Hs01101219_g1 | TaqMan® Gene Expression Assay | 4331182 |
| HMOX1 | Hs01110251 | TaqMan® Gene Expression Assay | 4331182 |

##### S3.3 Viability

We monitored cell viability from orbital shaking (88 RPM) and static culture conditions by incubating NOs in basal medium containing 1mg/mL resazurin for 3 hours at 37 °C (250 $\mu$L per well). After 3 hours, we took 100 $\mu$L per condition for colorimetric analysis at excitation 530/25 and emission 590/35 on a Synergy HT Biotek spectrophotometer. We tested each condition in quadruplicate. We used a set of triplicate media containing no cells as the negative control to determine the culture medium background.

##### S3.4 Quantification of NO area

We analyzed the images using ImageJ to quantify NO and necrotic areas. We calibrated images to a scale of 1.6005 pixels/$\mu$m. Furthermore, we manually outlined the whole NO boundary and individual dark regions using the region of interest (ROI) tools. We converted pixel areas to mm² units for area measurement using the formula:

$$Area\ in\ mm^2 = Pixel\ Area\ x \left(\frac{1}{1.6005}\right)^2 x10^{-6} \qquad \textbf{[S5]}$$

To quantify necrotic area formation across the NO, we summed the total dark regions and remaining tissue across slices and calculated the cumulative proportions as:

$$\%\ necrotic\ area = \frac{necrotic\ area\ (mm^2)}{total\ organoid\ area\ (mm^2)} \times 100 \qquad \textbf{[S6]}$$
